# LAMA2-related muscular dystrophy alters Schwann cell calcium homeostasis and increases peripheral nerve inflammation

**DOI:** 10.64898/2026.10.27.762325

**Authors:** John Wang, Mallika Khati, Molly Heron, Anushe Munir, Oluwaseun Akinyele, Marie A. Johnson, Gerard Vockley, Udai B. Pandey, Dwi U. Kemaladewi

## Abstract

LAMA2-related muscular dystrophy (LAMA2-RD) affects skeletal muscle and peripheral nerves, yet the consequences of LAMA2 deficiency on peripheral nerve integrity beyond early postnatal development remain incompletely understood. We investigated sciatic nerve architecture and molecular alterations across postnatal life in hypomorphic *Lama2^dy2J/dy2J^*mice. LAMA2 localizes around the abaxonal surface of myelinating Schwann cells in control nerves. LAMA2-deficient nerves exhibited patchy regions lacking myelinated axons from early postnatal stages into adulthood, including expanded Remak bundles along with axons completely lacking normal Schwann cell ensheathment. In adult nerves, myelin infolding and reduced myelin circularity revealed abnormalities in the architecture of myelin that had formed. Bulk sciatic nerve RNA sequencing identified altered calcium-signaling and immune-related transcriptional signatures. Complementary Fura Red measurements in primary nerve-derived Schwann cell cultures demonstrated decreased intracellular calcium in LAMA2-deficient samples, while ATP-evoked responses were preserved. Older LAMA2-deficient nerves also exhibited increased CD45^+^CD11b^+^ myeloid-cell infiltration. Together, these observations extend the characterization of LAMA2-associated neuropathy into adulthood and identify calcium homeostasis and neuroimmune responses as candidate processes relevant to persistent peripheral nerve pathology.

## Introduction

Biallelic pathogenic variants in the *LAMA2* gene cause LAMA2-related muscular dystrophy (LAMA2-RD) (1), a multisystemic disease that is one of the most common congenital muscular dystrophies (2–5). This disorder presents across a clinical spectrum, ranging from severe congenital muscular dystrophy to later-onset limb-girdle muscular dystrophy (6–9). *LAMA2* encodes laminin α2, which assembles with laminin β1 and γ1 to form laminin-211 (10, 11), a basement-membrane component that engages dystroglycan and integrin receptors (12–14). Although skeletal muscle degeneration is the most extensively studied manifestation, laminin-211 is also an important component of the peripheral nerve extracellular matrix (15, 16). Peripheral nerve involvement is therefore an integral, but comparatively understudied, feature of LAMA2-RD (17–19). Reduced nerve conduction velocities (19–22) and abnormal myelin morphology (17, 20, 23, 24) have been described in individuals with LAMA2-RD, and experimental models demonstrate abnormal Schwann cell radial sorting and bundles of incompletely ensheathed axons (18, 25, 26). These abnormalities may contribute to motor impairment independently of primary muscle pathology. This distinction is relevant to emerging treatments directed predominantly toward skeletal muscle (27, 28), because improvement in muscle function may not fully address persistent nerve pathology.

Much of the experimental literature has focused on early myelination (18, 25), in part because severe *Lama2*-deficient mouse models have limited survival (29–32). The hypomorphic *Lama2^dy2J/dy2J^* model, with a near-normal lifespan (33, 34), provides an opportunity to investigate whether peripheral nerve abnormalities persist and whether additional changes in myelin architecture emerge in adulthood. It also permits investigation of molecular pathways potentially involved in Schwann cell function and peripheral nerve maintenance.

Here, we characterized the distribution of LAMA2 in sciatic nerve, examined myelin architecture and axonal ensheathment across postnatal ages, and profiled transcriptional changes in LAMA2-deficient nerves. Guided by the RNA-sequencing results, we assessed intracellular calcium levels in primary nerve-derived cultures and myeloid-cell abundance in nerve tissue. Our findings connect persistent structural pathology with calcium-related and neuroimmune alterations, providing a framework for investigating the maintenance of peripheral nerve integrity in LAMA2-RD.

## Results

### LAMA2 localizes to the abaxonal surface of myelinating Schwann cells

Laminin α2 is a major component of the Schwann cell basement membrane (15, 16), and its spatial distribution relative to compact myelin and the axon is important for understanding how LAMA2 deficiency disrupts peripheral nerve development. In particular, localization at the abaxonal Schwann cell surface would place laminin-211 near extracellular matrix receptors implicated in radial sorting and myelin organization (15, 35–38), whereas localization at the adaxonal interface would suggest a different relationship with axon–Schwann cell contact (Figure 1, A). We therefore examined the distribution of LAMA2 relative to myelinating Schwann cells expressing myelin basic protein (MBP) (39) and axons expressing βIII-tubulin (40) in wild-type sciatic nerve cross-sections at 1, 2, and 6 weeks of age (Figure 1, B). At all three ages, LAMA2 immunoreactivity formed a ring external to MBP-positive compact myelin and surrounded βIII-tubulin-positive axons. The signal was enriched at the abaxonal surface of myelinating Schwann cells rather than at the axon-facing interface. This pattern was also observed in 1-week-old animals, where the axon remained near the periphery of the myelinating Schwann cell, which likely represents the initial stages of myelination (Figure 1, B arrow). In 25-week-old adult animals, LAMA2 was also detected at the outer circumference of Remak bundles – unmyelinated, small-diameter axon groupings surrounded by the cytoplasmic protrusions of a Remak Schwann cell (41, 42) (Supplemental Figure 1). Secondary-antibody-only controls showed minimal background signal (Supplemental Figure 2). These observations define the anatomical context for investigating extracellular matrix–dependent Schwann cell signaling.

**Figure 1:**
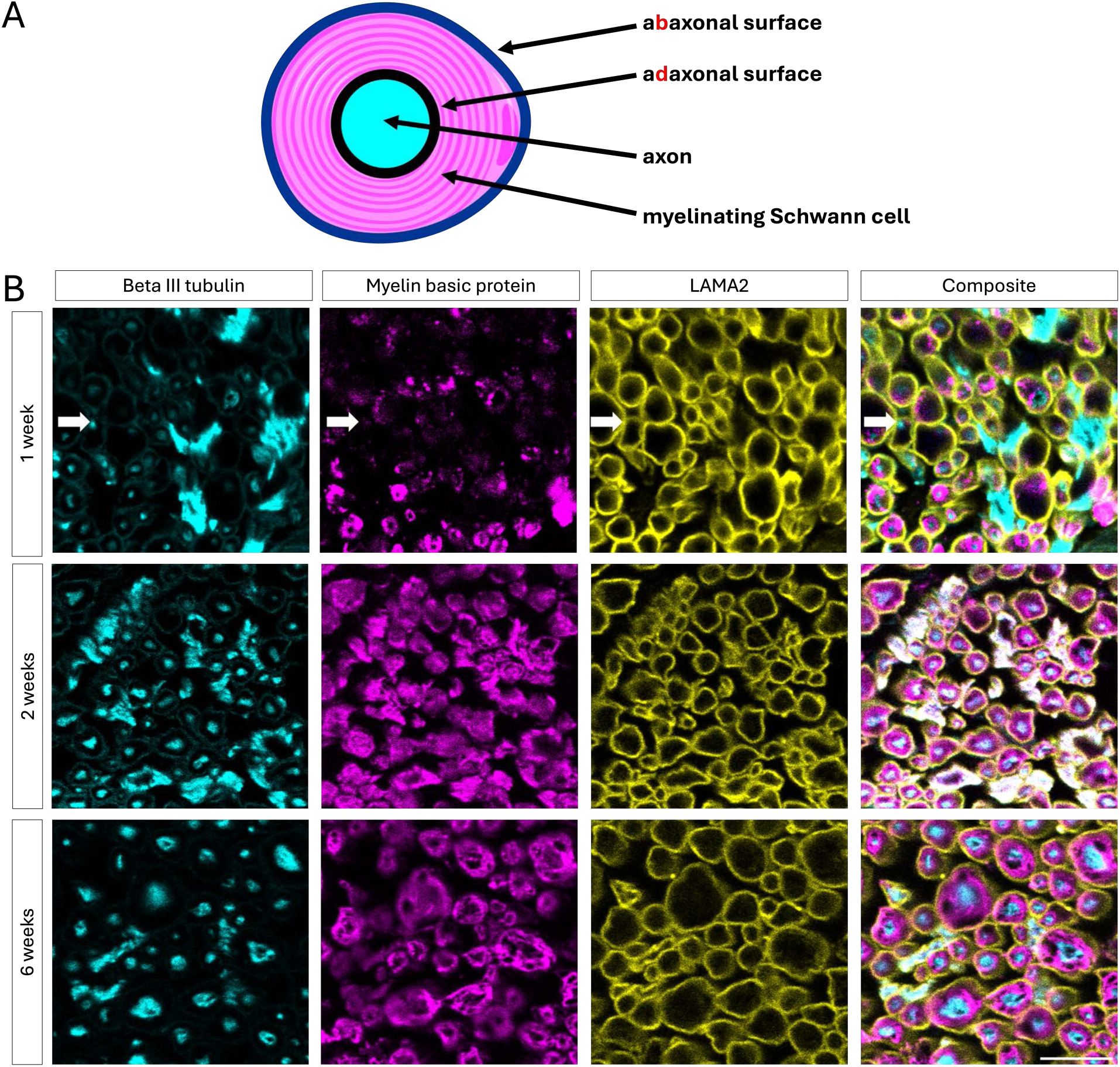
LAMA2 protein localizes to the Schwann cell abaxonal surface. A) Schematic of abaxonal and adaxonal surfaces of myelinating Schwann cells. B) Representative sciatic nerve cross-sections from wild-type C57BL/6J mice of the indicated ages immunostained against axons (β-III tubulin), myelinating Schwann cells (myelin basic protein), and LAMA2. Arrow indicates axon at the periphery of a LAMA2-positive circular region. LAMA2 immunoreactivity outlines myelinated fibers on the abaxonal side of the myelin sheath. Sciatic nerves from 3-5 mice of each age were assessed. Scale bar = 10 µm.

### LAMA2-deficient nerves show persistent amyelinated regions and age-associated myelin abnormalities

Because laminin-211 is required for normal Schwann cell radial sorting (18, 25, 26), we asked whether defects established during development remain apparent in older nerves and whether myelin that successfully forms retains normal architecture. The longer-lived hypomorphic model enables assessment of adult stages that are difficult to study in severe LAMA2-null models, which do not survive past the juvenile and early adult stages (29–32). We evaluated sciatic nerve morphology in wild-type and Lama2^dy2J/dy2J^ mice across postnatal development and adulthood. Toluidine blue–stained cross-sections revealed patchy regions lacking myelinated axons as early as 2 weeks in LAMA2-deficient nerves (Figure 2, A). These regions remained apparent at subsequent ages. At 25 and 52 weeks, the fraction of nerve cross-sectional area lacking myelinated axons was greater in LAMA2-deficient nerves than in wild-type controls (Figure 2, B), demonstrating that the abnormal distribution of myelinated fibers persists into adulthood. High-magnification electron microscopy revealed these regions to be a combination of enlarged Remak bundles and bundles of unmyelinated axons that are also not associated with any Remak Schwann cells (Supplemental Figure 3).

**Figure 2:**
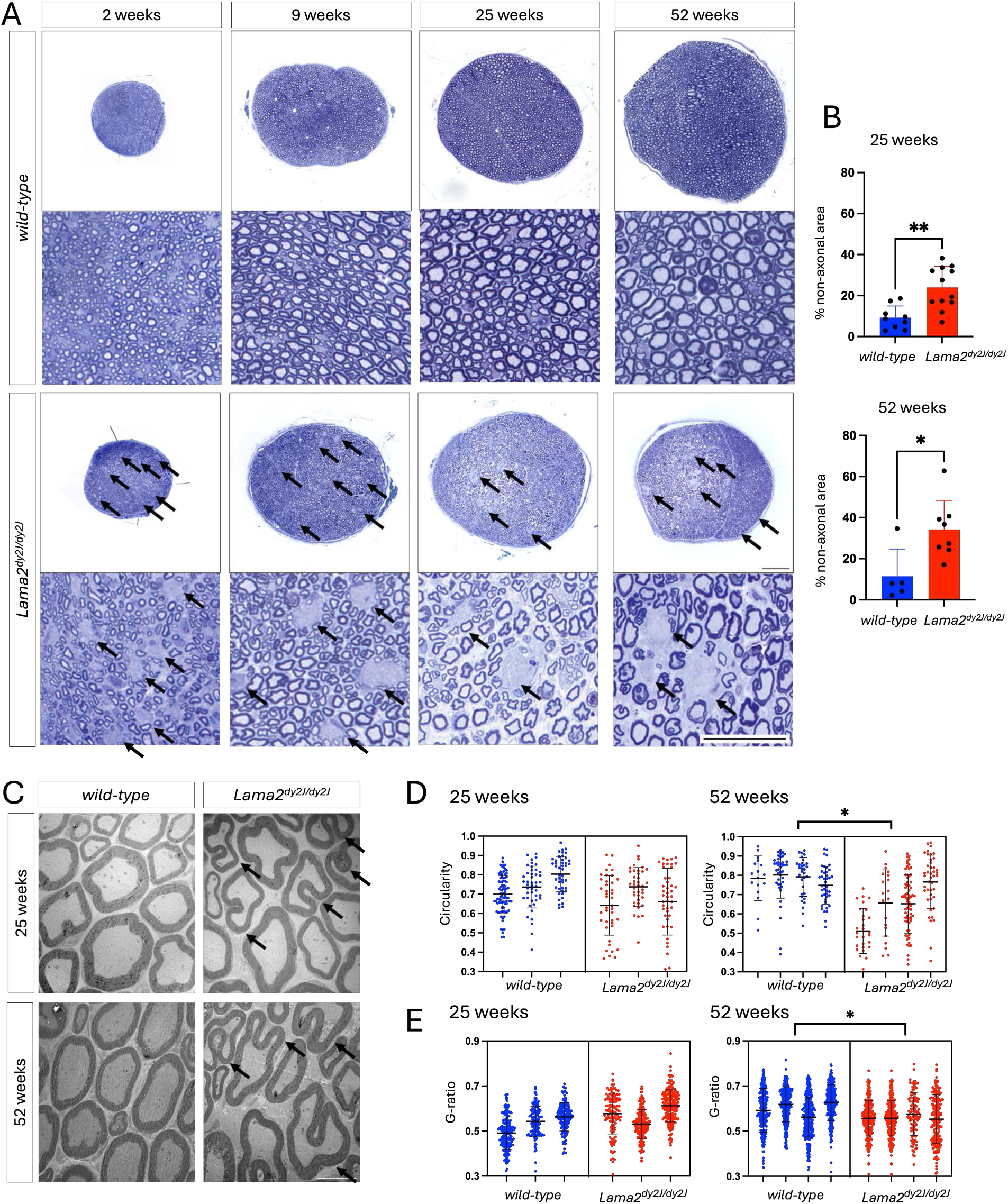
LAMA2-deficient sciatic nerves exhibit persistent regions lacking myelinated axons and abnormal myelin morphology in adulthood. A) Toluidine-blue staining of sciatic nerve cross-sections from mice of the indicated genotypes and ages. Top panels are overviews of the entire nerve, and bottom panels are a zoomed-in view of a portion of the nerve. Arrows indicate regions lacking myelinated axons. Scale bar = 100 µm for top panels, 50 µm for bottom panels. B) Quantification of percentage of total nerve cross-sectional area that lacks myelinated axons for the indicated genotypes and ages. Each point represents an individual mouse. Bar heights represent the mean, and error bars represent standard deviation (SD). Two-tailed Welch’s t test was performed. * p < 0.05, ** p < 0.01. C) Transmission electron micrographs of sciatic nerve cross-sections from mice of the indicated genotypes and ages taken at 5000× magnification. Arrows indicate myelinated axons with increased myelin infolding. Scale bar = 5 µm. D) Quantification of myelin circularity for sciatic nerves from mice of the indicated ages and genotypes. Each column represents an individual mouse, and each dot represents an individual myelinated axon. Horizontal line indicates the mean, and error bars represent SD. Two-tailed nested t test was performed. * p < 0.05. E) Quantification of g-ratio (a measure of myelin thickness – lower ratios indicate thicker myelin) for sciatic nerves from mice of the indicated ages and genotypes. Each column represents an individual mouse, and each dot represents an individual myelinated axon. Horizontal line indicates the mean, and error bars represent SD. Two-tailed nested t test was performed. * p < 0.05.

Electron microscopy further revealed abnormalities in the myelin formed around remaining myelinated axons. Myelin infoldings were apparent at 25 and 52 weeks in LAMA2-deficient nerves (Figure 2, C), and quantitative measurements showed reduced myelin circularity at 52 weeks (Figure 2, D). The g-ratio was also lower at 52 weeks (Figure 2, E). In the setting of irregular myelin contours and infoldings, this finding is most informative as part of the broader architectural phenotype rather than as evidence of thicker myelin.

These cross-sectional observations establish differences in peripheral nerve myelination across sampled ages at the population level. Overall, LAMA2 deficiency is associated with persistent radial-sorting/ensheathment abnormalities as well as alterations in the organization of adult myelin.

### Bulk RNA sequencing identifies signatures of altered calcium signaling and upregulated inflammatory pathways

Structural analysis established the persistence of abnormal ensheathment and adult myelin morphology but did not identify the cellular programs associated with these phenotypes. Because laminin-211 interacts with cell-surface receptors, such as integrin and dystroglycan, which are capable of influencing intracellular signaling, we asked which transcriptional pathways differ in LAMA2-deficient sciatic nerves. To investigate molecular changes associated with these structural abnormalities, we performed bulk RNA sequencing of sciatic nerves and evaluated differentially expressed genes using Qiagen’s Ingenuity Pathway Analysis (IPA; Figure 3, A). At 25 weeks, 288 genes were upregulated and 440 were downregulated (see Methods for differential-expression criteria). Calcium signaling was among the most statistically significantly perturbed pathways, with a negative predicted activation score, while immune-related programs, including neutrophil degranulation, showed positive predicted activation scores (Figure 3, B, D, F, G). At 52 weeks, calcium signaling continued to show a negative predicted activation score and multiple immune-related pathways were prominent (Figure 3, C, E-G).

**Figure 3:**
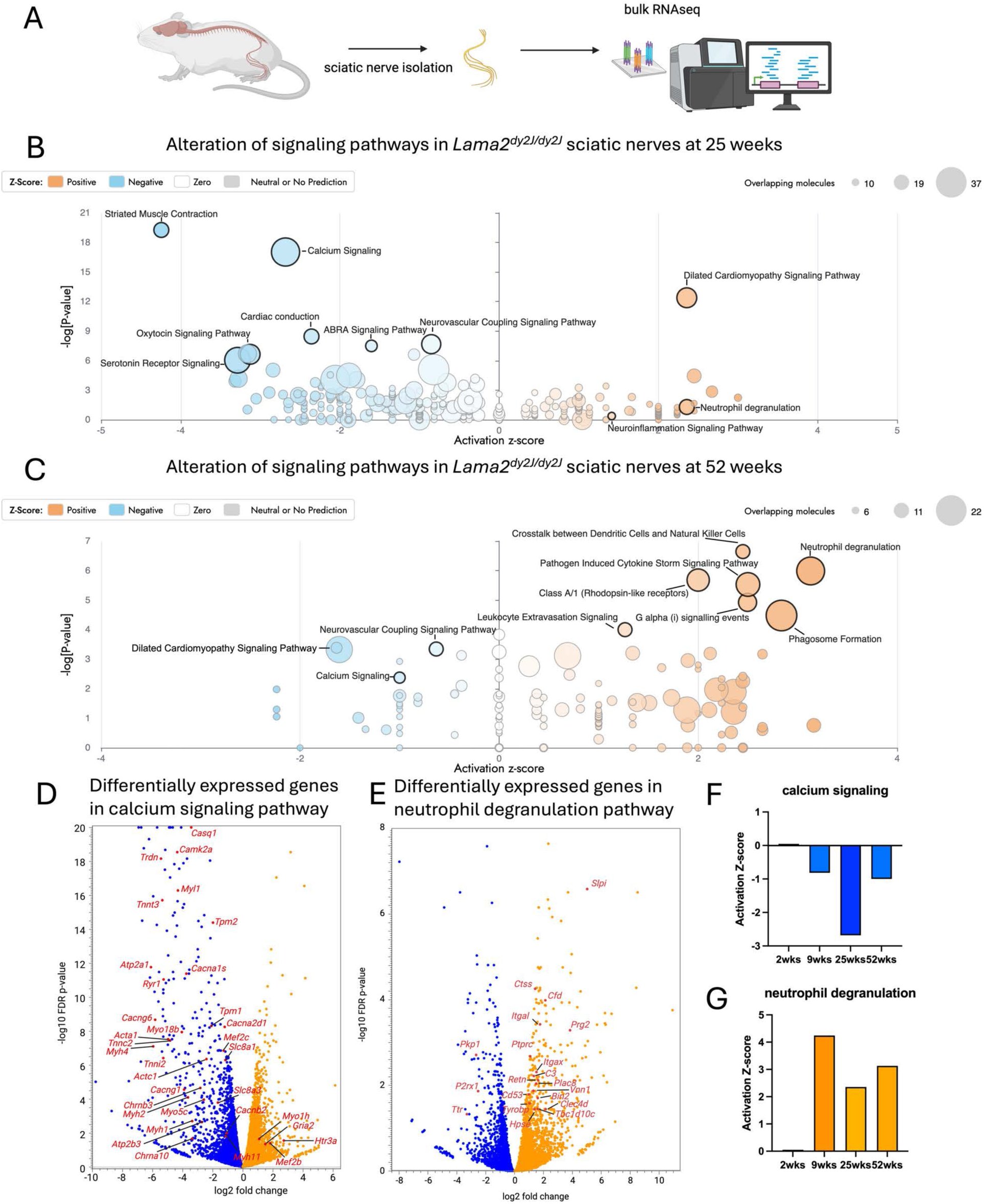
Bulk sciatic nerve RNA sequencing identifies altered calcium-related and immune-related transcriptional signatures in LAMA2-deficient nerves. A) Schematic of sciatic nerve isolation and bulk RNA sequencing. Nerves from individual mice were processed and sequenced separately. Pathway analysis was performed by pooling mice of the same genotype together. Nerves from five mice of each genotype at 25 weeks and four mice of each genotype at 52 weeks were analyzed. B) Volcano plot of signaling pathways predicted to be altered based on RNA sequencing data at 25 weeks. X axis represents activation z-score, with negative values indicated by blue bubbles and positive values indicated by orange bubbles. Y axis represents statistical significance, with high values representing increased statistical significance. The size of each bubble corresponds to the number of differentially expressed genes mapped to that pathway. C) Volcano plot of signaling pathways predicted to be altered based on RNA sequencing data at 52 weeks. Plot is labeled in the same manner as panel B. D) Volcano plot of differentially expressed genes in the calcium signaling pathway (red dots) at 25 weeks overlaid on all differentially expressed genes. X axis represents log_2_ fold change in expression and Y axis represents statistical significance, with higher values indicating increased statistical significance. E) Volcano plot of differentially expressed genes in the neutrophil degranulation pathway at 52 weeks. Plot is labeled in the same manner as panel D. F) Predicted activation z-score of the calcium signaling pathway at the indicated timepoints. Earlier timepoint data was based on analysis of nerves from 6 mice of each genotype at 2 weeks and 4 mice of each genotype at 9 weeks. G) Predicted activation z-score of the neutrophil degranulation pathway at the indicated timepoints.

Across adult timepoints, the calcium-signaling signature and immune-associated signatures were consistently represented in the pathway analyses (Figure 3, F and G; Supplemental Figure 4). Analysis of sciatic nerves from a separate severe Lama2^dyW/dyW^ model (30) at 2 weeks also identified calcium-related and immune-associated pathway changes (Supplemental Figure 5). Together, these analyses identified recurring calcium-signaling and inflammatory transcriptional signatures across multiple disease stages and LAMA2-deficient mouse models.

To further characterize the molecular changes underlying these pathway-level alterations, we examined differentially expressed genes associated with calcium signaling and inflammatory responses. Genes contributing to the calcium-signaling signature included *Cacna1s, Ryr1, Trdn, Casq1, Atp2a1,* and other calcium-associated transcripts (Figure 3, D; Supplemental Figure 6). These genes encode proteins involved in calcium transport, intracellular calcium storage and release, and calcium-dependent cellular processes (43–47). Similarly, genes contributing to the neutrophil degranulation pathway included *Cfd, C3, Itgal, Itgax,* and other transcripts that participate in innate immune responses and/or leukocyte adhesion (Figure 3, E) (48–51).

Together, these transcriptomic findings identified calcium signaling and inflammation as two prominent molecular pathways altered in LAMA2-deficient peripheral nerves. To complement these findings with cellular and histological evidence, we next investigated intracellular calcium homeostasis in primary nerve-derived cultures and assessed immune-cell accumulation in sciatic nerves.

### LAMA2 deficiency is associated with a lower basal Fura Red ratio in primary Schwann cell cultures

The calcium-related transcriptomic signature prompted us to test whether an independent functional readout also differed in cells derived from LAMA2-deficient nerves. Because intracellular calcium participates in Schwann cell responses to extracellular cues (52, 53), we examined both basal conditions and the response to ATP, a known stimulator of Schwann cell intracellular calcium release (54, 55). We assessed calcium-dependent fluorescence using the ratiometric indicator Fura Red (56, 57) in primary cultures derived from wild-type and *Lama2^dy2J/dy2J^* sciatic nerves (Figure 4, A). The cultures included S100B^+^SOX10^+^ Schwann cells (58–60) and other nerve-derived cell types (Supplemental Figure 7). Flow-cytometric analysis excluded debris, doublets, nonviable cells, and Thy1.2-positive fibroblast cells (61, 62) before evaluating Fura Red ratios (Supplemental Figure 8).

**Figure 4:**
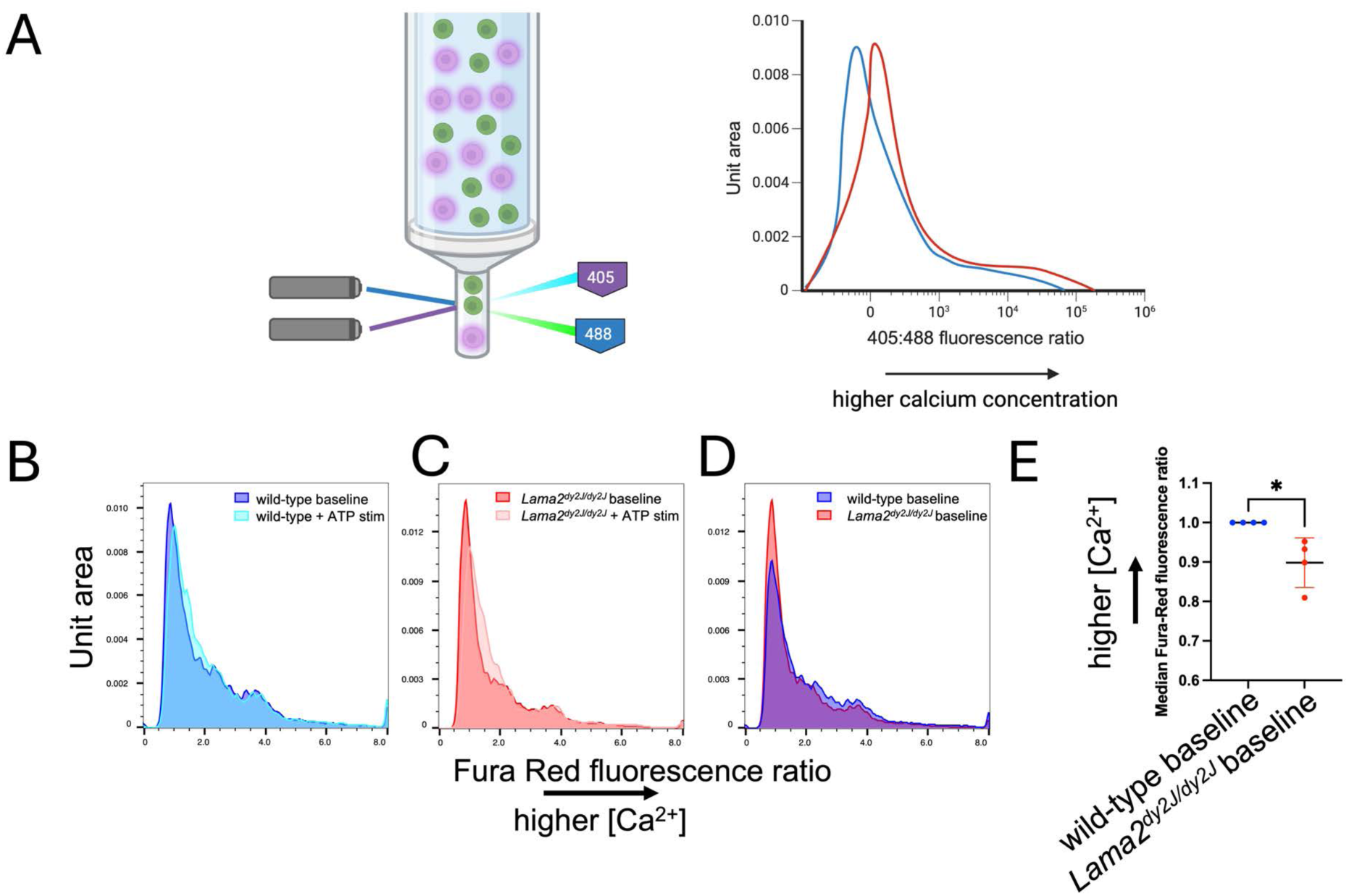
LAMA2 deficiency is associated with a decreased basal Fura Red fluorescence ratio in nerve-derived primary Schwann cell cultures. A) Schematic of intracellular calcium measurements using flow cytometry. Cells are loaded with Fura Red and excited with 405-nm and 488-nm lasers. The 405:488 ratio increases with intracellular calcium concentration. B-D) Representative 405:488 ratio histograms of a single experiment with primary Schwann cells of the indicated genotype with and without ATP stimulation. Plots are normalized so that the total area under the curve equals 1.0. Higher ratios correspond to higher intracellular calcium concentration. E) Plot of normalized median Fura Red 405:488 fluorescence ratio across four biological replicates. Each dot represents an experimental replicate performed on a separate day from a separate batch of primary Schwann cells. Line represents the mean, and error bars represent SD. Two-tailed Welch’s t test was performed. * p < 0.05.

ATP stimulation shifted the Fura Red excitation-ratio distributions in both genotypes (Figure 4, B and C), consistent with retained stimulus-responsive calcium signaling. In unstimulated cultures, the basal Fura Red ratio was lower in LAMA2-deficient samples than in wild-type controls (Figure 4, D), with a corresponding difference across multiple biological replicates summarized in Figure 4, E. These results identify an alteration in basal Schwann cell intracellular calcium levels associated with LAMA2 deficiency.

### Older LAMA2-deficient nerves contain increased numbers of CD45^+^CD11b^+^ myeloid cells

The immune-related RNA-sequencing signatures raised a complementary question: whether these transcriptional changes were accompanied by an observable difference in immune cells within peripheral nerve tissue. To assess this independently of pathway inference, we stained for immune-cell markers in sciatic nerve sections. Because many of the upregulated RNA-sequencing pathways were related to innate immunity, and because macrophage infiltration has been reported in other peripheral neuropathies, including Charcot-Marie-Tooth disease and diabetic peripheral neuropathy (63–67), we used CD45 to identify leukocytes, CD11b to identify myeloid cells, and CD68 to identify macrophages (68–70). At 9 weeks, CD45^+^CD11b^+^ myeloid cell abundance was not significantly different between wild-type and LAMA2-deficient nerves (Figure 5, A and B). By 52 weeks, LAMA2-deficient nerves contained more CD45^+^CD11b^+^ cells than controls (Figure 5, C and D). A subset of CD11b^+^ cells co-expressed CD68 (Figure 5, C), consistent with the presence of macrophages, whereas the majority were CD68-negative, suggesting that a different myeloid population predominated. These histologic observations complement the immune-associated transcriptional signature and reveal increased myeloid-cell abundance in older LAMA2-deficient peripheral nerves.

**Figure 5:**
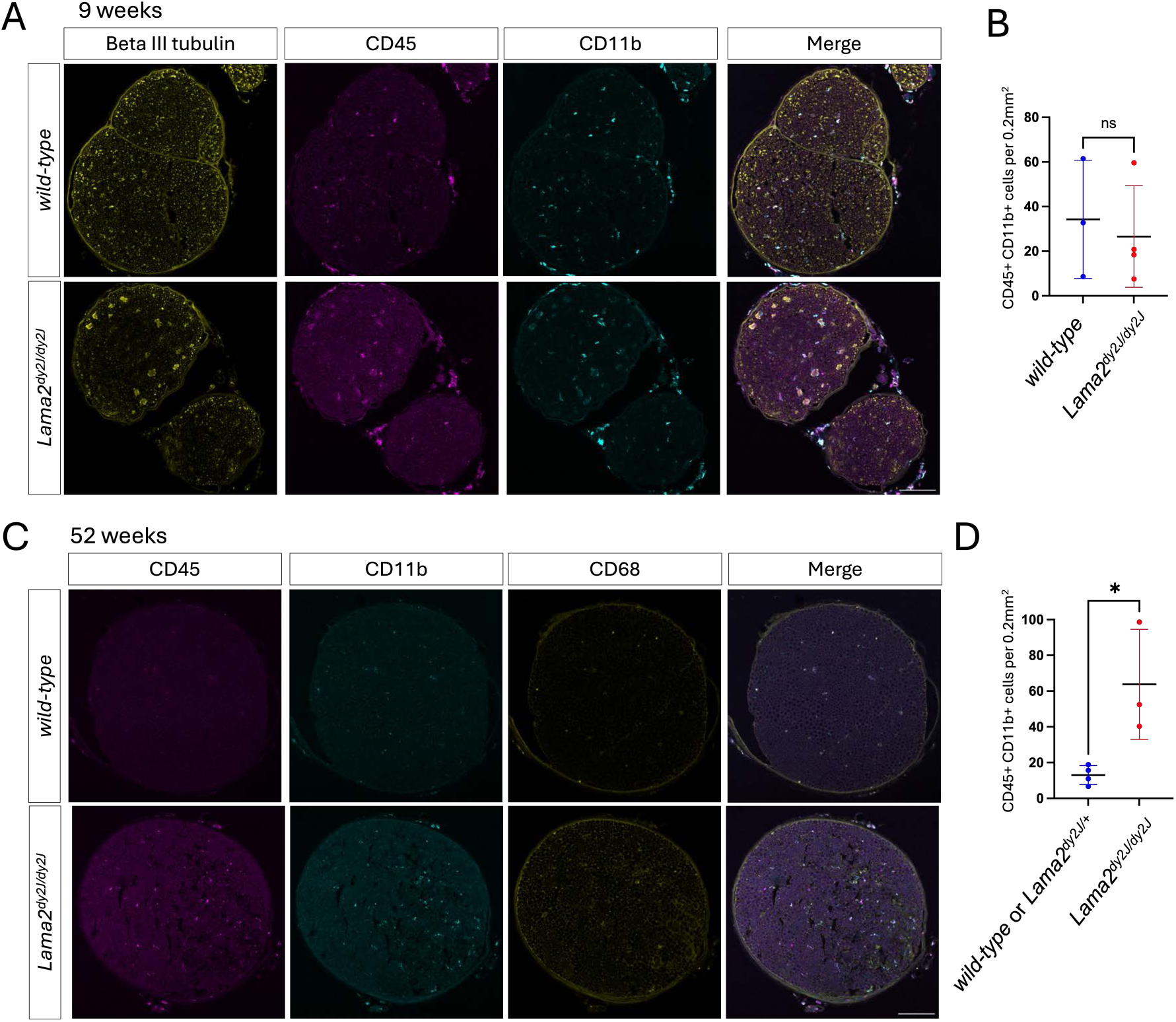
Increased myeloid-marker-positive cells in sciatic nerves of older LAMA2-deficient mice. A) Representative image of sciatic nerves from 9-week-old mice of the indicated genotypes immunostained against the indicated antigens. Scale bar = 100 µm. B) Quantification of CD45^+^CD11b^+^ myeloid cells in nerves of the indicated genotype at 9 weeks. Each dot represents an individual nerve. Horizontal lines represent the mean. Error bars represent SD. Two-tailed Welch’s t test was performed. NS = not statistically significant. C) Representative image of sciatic nerves from 52-week-old mice of the indicated genotypes immunostained against the indicated antigens. Scale bar = 100 µm. D) Quantification of CD45^+^CD11b^+^ myeloid cells in nerves of the indicated genotype at 52 weeks. Each dot represents an individual nerve. Horizontal lines represent the mean. Error bars represent SD. Two-tailed Welch’s t test was performed. * p < 0.05.

## Discussion

Our study extends the characterization of LAMA2-associated peripheral neuropathy beyond the early stages of radial sorting and myelination. Three findings are particularly informative. First, patchy regions lacking normally myelinated axons persist into adulthood and contain both enlarged Remak bundles and axonal bundles lacking Schwann cell ensheathment. Second, the myelin that does form exhibits architectural abnormalities in adult LAMA2-deficient nerves, including infolding and altered circularity. Third, molecular and cellular analyses identify calcium-related alterations and a myeloid-cell infiltrate as features of the peripheral nerve phenotype. Together, these findings support a broader role for laminin-211 in maintaining the homeostasis of peripheral nerves throughout postnatal life.

LAMA2 localization provides a useful anatomical context for these observations. The enrichment of laminin α2 external to compact myelin is consistent with its established position within the Schwann cell basement membrane and its interactions with integrin and dystroglycan receptors (15, 35, 36, 71). Such contacts are positioned to influence Schwann cell morphology, adhesion, and responses to the extracellular environment. The localization also raises the possibility that laminin-211 contributes to mechanical stability in the peripheral nerve, analogous to its structural role in muscle basement membranes. During ordinary limb movement, peripheral nerves experience stretch and deformation (72, 73); altered matrix support due to loss of laminin-211 could conceivably increase vulnerability to myelin disruption, which has been shown in other alterations that disrupt the basal lamina (74, 75). Although not addressed by the present experiments, direct tests of this mechanical support hypothesis may elucidate additional roles of LAMA2 in maintaining peripheral nerve homeostasis.

The structural findings suggest that LAMA2 deficiency affects more than the initiation of myelination. The naked axonal bundles are consistent with a persistent radial-sorting and ensheathment defect (76, 77), whereas infolding and loss of circularity indicate abnormal morphology of the myelin present in adult nerves, which may reflect a defect in myelin formation or maintenance (78–80). The lower g-ratio at 52 weeks should be considered in this morphological context: redundant or infolded membrane can alter apparent sheath geometry without implying more effective myelination. Similar myelin abnormalities with increased infolding (36) or deficits in radial sorting (77) in models disrupting Schwann cell integrin- and dystroglycan-dependent interactions make extracellular matrix–receptor signaling a compelling direction for future studies. Conditional approaches that separate developmental from adult requirements for LAMA2 (e.g., using Schwann cell-specific or inducible *Lama2* deletion (81)) would help determine whether the later phenotype reflects impaired myelin maintenance, cumulative effects of an early myelination defect, or both.

Another important observation is the association between LAMA2 deficiency and altered calcium homeostasis. The bulk RNA-sequencing data identified a calcium-related transcriptional signature, and the Fura Red experiments independently detected a difference in basal calcium-dependent fluorescence in LAMA2-deficient nerve-derived cultures. Notably, ATP-evoked responses remained detectable in both genotypes, indicating that the basal difference does not correspond to a complete loss of stimulus responsiveness, at least to supramaximal ATP stimulation. These observations raise the possibility that the extracellular matrix environment influences resting calcium regulation in peripheral nerve cells.

There is biological precedent for this hypothesis. Laminin-binding receptors, including integrins and dystroglycan, participate in signaling networks that can intersect with focal adhesion kinase, phosphatidylinositol 3-kinase, and calcium-regulatory pathways (82–84). Calcium dynamics, in turn, influence Schwann cell development, differentiation, axon interactions, and responses to injury. Prior studies of peripheral neuropathies have illustrated that both excessive and insufficient calcium signaling may disrupt cellular homeostasis (53, 85–87), emphasizing the importance of appropriate regulation rather than simple activation or inhibition. In this context, the altered basal Fura Red ratio suggests a candidate cellular consequence of laminin-211 deficiency that warrants mechanistic investigation. Identifying the relevant calcium stores, channels, and receptor-linked pathways, and testing their effects on myelination, would help establish whether this association contributes causally to disease.

The increased abundance of CD45^+^CD11b^+^ cells in older LAMA2-deficient nerves adds a neuroimmune dimension to the phenotype. Peripheral nerve macrophages and other myeloid cells can participate in debris clearance, tissue remodeling, and repair (88–90), but persistent immune activation can also worsen chronic neuropathology (63–67). The present observations do not determine whether these cells are primarily protective, pathogenic, or neutral bystanders. Resolving their functional state and effect on disease pathology will be important before considering immune modulation as a therapeutic strategy.

These findings are relevant to therapeutic development in LAMA2-RD. Many natural history studies of LAMA2-RD do not assess for peripheral neuropathy (91–95), and some therapeutics in development do not address it (27). Interventions that primarily improve skeletal muscle pathology are unlikely to correct peripheral nerve defects that originate in development or persist in adulthood (26, 96, 97). Assessing nerve architecture and function alongside muscle outcomes could provide a more complete understanding of disease natural history and a more holistic evaluation of treatment benefit. The calcium and neuroimmune observations identify promising avenues for mechanistic work that may eventually identify novel therapeutic targets.

In summary, LAMA2 deficiency is associated with persistent axonal ensheathment defects, abnormal adult myelin architecture, altered basal calcium-dependent signaling in nerve-derived primary Schwann cell cultures, and increased myeloid-cell abundance in older peripheral nerves. These findings broaden our understanding of LAMA2-associated neuropathy and motivate studies of how extracellular matrix support, calcium regulation, and neuroimmune responses contribute to peripheral nerve integrity.

## Methods

### Sex as a biological variable

Male and female mice were included in approximately equal proportions across experimental groups. Sex was considered a biological variable in the experimental design; however, the studies were not powered to detect sex-specific effects, and data from both sexes were pooled for analysis. Further studies are needed to determine whether peripheral nerve pathology in LAMA2-RD differs by sex.

### Mice

Lama2^dyW^ (Strain #:013786) and Lama2^dy2J^ (Strain #:000524) mice were initially obtained from The Jackson Laboratory and colonies were maintained by intercrossing heterozygous mice within each strain. Genotyping was performed either by Transnetyx or in-house using PCR-based assays of DNA from ear tissue. Both male and female animals were used for experiments in approximately equal proportions, with ages ranging from 1 week to 52 weeks.

### Genotyping

For *Lama2^dyW^* genotyping, multiplex PCR used primers 5′-ACTGCCCTTTCTCACCCACCCTT-3′, 5′-GTTGATGCGCTTGGGACTGTCG-3′, and 5′-CGACAGTATCGGCCTCAGGAAG-3′ with Platinum Multiplex PCR Master Mix (Applied Biosystems). Cycling conditions were 95°C for 30 seconds, 68°C for 60 seconds, and 72°C for 30 seconds for 35 cycles. Products were resolved on 2% agarose gels (wild-type, 246 bp; *Lama2^dyW^*, 446 bp).

For *Lama2^dy2J^* genotyping, PCR used primers 5′-TTGCTTCGAATGCACTCATC-3′ and 5′-TGGATGTATATGGGACACTAATG-3′ with PCR SuperMix (Invitrogen). Cycling conditions were 94°C for 15 seconds, 52°C for 30 seconds, and 72°C for 30 seconds for 40 cycles. PCR products were digested with NdeI (New England Biolabs, R0111S) at 37°C for 2 hours and resolved on 2% agarose gels (wild-type, 347 bp; *Lama2^dy2J^*, 202 bp and 145 bp).

### Sciatic nerve immunofluorescence

Mice were euthanized by CO_2_ inhalation followed by cervical dislocation. Sciatic nerves were isolated, fixed in 4% paraformaldehyde at 4°C overnight, washed in PBS, cryoprotected sequentially in 15% and 30% sucrose, embedded in OCT compound, and cryosectioned at 10 μm. Sections were permeabilized in 0.2% Triton X-100 in PBS for 5 minutes, blocked in 2% BSA plus 0.1% Tween-20 in PBS, and incubated with primary antibodies for 1 hour at room temperature or overnight at 4°C. After washing, sections were incubated with secondary antibodies and DAPI for 1 hour at room temperature, washed, and mounted in ProLong Gold antifade mountant.

Primary antibodies were rabbit anti-MBP (Abcam, ab40390), rat anti-LAMA2 (Abcam, ab11576), mouse anti–βIII-tubulin/TUJ1 (BioLegend, 801213), rat anti-CD11b (Cell Signaling Technology, 46512S), rabbit anti-CD45 (Cell Signaling Technology, 70257S), and mouse anti-CD68 (NeoBiotechnologies, 968-MSM2-P0). Secondary antibodies were goat anti-rabbit Alexa Fluor 594 (Invitrogen, A32740), goat anti-rat Cy5 (Invitrogen, A10525), goat anti-mouse Alexa Fluor 488 (Invitrogen, A32723), donkey anti-rabbit Alexa Fluor 488 (Thermo Fisher Scientific, A21206), donkey anti-rat Alexa Fluor 568 (Thermo Fisher Scientific, A78946), and donkey anti-mouse Alexa Fluor 647 (Thermo Fisher Scientific, A31571).

### Confocal microscopy

High-resolution LAMA2 co-stains were imaged on a Leica Stellaris 5 inverted confocal microscope with a white-light laser and Power HyD S detectors using a 63×/1.4 NA oil-immersion objective. Images of myelinated fibers were acquired at 2048 × 2048 pixels with an XY pixel size of approximately 90.1 nm. Images of Remak bundles were acquired at 2048 × 2048 pixels at 4× zoom with an XY pixel size of approximately 22.5 nm. Pinhole was set to 1 Airy unit and line averaging of 4 was employed. DAPI/594 and 488/Cy5 channels were acquired in sequential tracks, with tracks alternating with each line. Immune-marker staining was imaged on the same microscope using a 20×/0.75 NA objective at 1024 × 1024 pixels with a 1-Airy-unit pinhole and line averaging of 4. Secondary-antibody-only controls were processed in the same batch and imaged using the same acquisition settings as corresponding experimental samples.

### RNA sequencing and pathway analysis

Sciatic nerves were snap-frozen in liquid nitrogen or immediately immersed in RNAlater. Tissue was homogenized in TRIzol using 1.4-mm ceramic beads and a Bullet Blender tissue homogenizer (Next Advance), and total RNA was isolated according to the manufacturer’s protocol.

Stranded total RNA libraries were generated with the Illumina Stranded Total Library Prep kit (Illumina: 20040529) according to the manufacturer’s instructions. Briefly, 20-50 ng of input RNA was used for each sample. Following adapter ligation, 12-15 cycles of indexing PCR were completed, using Illumina RNA UD Indexes. Library quantification and assessment were done using a Qubit FLEX fluorometer and an Agilent TapeStation 4150/Fragment Analyzer 5300. Libraries were normalized and pooled to 2 nM by calculating the concentration based on the fragment size (base pairs) and the concentration (ng/µL) of the libraries. Sequencing was performed on an Illumina NextSeq 2000. The pooled library was loaded at 750 pM and sequencing was carried out with read lengths of 2 × 101 bp, with a target of 40 million reads per sample.

Sequencing data were demultiplexed by the on-board Illumina DRAGEN FASTQ Generation software. Reads were aligned to *Mus musculus* GRCm39 using QIAGEN CLC Genomics Workbench v24. Parameters: mismatch cost = 2, insertion cost = 3, deletion cost = 3, length fraction = 0.8, similarity fraction = 0.8, global alignment = no, auto-detect paired distances = yes, maximum number of hits for a read = 10, strand setting = both, library type setting = bulk, count paired reads as two = no, ignore broken pairs = yes, expression value = total counts, calculate expression for genes without transcripts = no.

Differential expression was analyzed using the CLC two-group differential-expression workflow. For pathway analysis, differentially expressed genes were filtered using |log2 fold change| > 1 and FDR-adjusted P < 0.05 and loaded into Qiagen Ingenuity Pathway Analysis (IPA) software. Core analysis was performed in IPA, and activation z-scores were used to summarize predicted pathway activation or inhibition.

### Toluidine blue staining and transmission electron microscopy

Sciatic nerves were fixed in Karnovsky fixative (2.5% glutaraldehyde and 2% paraformaldehyde in PBS), rinsed, post-fixed in 1% osmium tetroxide with 1% potassium ferricyanide, dehydrated through graded ethanol and propylene oxide, and embedded in Poly/Bed 812. Semithin 300-nm sections were stained with 0.5% toluidine blue in 1% sodium borate. Ultrathin 65-nm sections were stained with UranyLess and Reynolds lead citrate and imaged on a JEOL 1400 Plus transmission electron microscope equipped with an AMT 2k camera.

### Image acquisition of toluidine blue-stained nerve sections and quantification of non-myelinated area

Slides were scanned using a Leica Aperio CS2 Slide Scanner at 40× magnification. Regions of interest (ROIs) corresponding to nerve sections were manually selected using Leica Aperio ImageScope software and converted to TIFF files. TIFF files were thresholded in Fiji to identify myelinated area and convert the values of those corresponding pixels to 255, while assigning all other pixels a value of 0. This was done by converting the original TIFF image to an 8-bit image, running “Enhance Local Contrast” (block size = 127, histogram = 256, maximum = 3) followed by “Auto Local Threshold” with the Phansalkar method (radius = 15), and then filling holes with the “Fill Holes” command. The nerve boundaries were outlined manually, and the total number of pixels was measured in addition to the total pixel intensity. Total pixel intensity was divided by 255 to calculate the number of pixels corresponding to myelinated area, and this was divided by the total number of pixels within the nerve to obtain the percent area corresponding to myelinated axons (axon + surrounding myelin sheath). This was subtracted from 100% to obtain the “% non-axonal area” shown in Figure 2, B.

### G-ratio and circularity analysis

For g-ratio analysis, images were thresholded in Fiji using the Moments auto-threshold algorithm and analyzed with MyelTracer (98). Axon and myelin contours were manually corrected when necessary. G-ratio values were exported for statistical analysis. For circularity, the outer myelin border was manually traced in Fiji and circularity was calculated as 4π × area/perimeter^2^.

### Immune-cell quantification

CD45^+^CD11b^+^ cells were quantified in Fiji and divided by the cross-sectional area of the nerve. This number was then scaled to be equivalent to a 0.2 mm^2^ cross-sectional area, which roughly corresponds to the average area of all nerves. A CD45-positive mask was generated by thresholding and applied to the CD11b channel; double-positive objects were then quantified using the 3D Object Counter v2.0 plugin with edge objects excluded.

### Primary nerve-derived cell culture

Sciatic nerves from 9- to 14-week-old wild-type or *Lama2^dy2J/dy2J^*mice were placed in ice-cold DMEM (Gibco) following isolation from the mouse. Nerve segments were subsequently transferred to tissue culture plates and underwent in vitro predegeneration for 6–7 days at 37°C in DMEM (Gibco) supplemented with 10% FBS (Gibco), GlutaMAX (Gibco), penicillin/streptomycin (Gibco), 10 nM recombinant neuregulin EGF domain (R&D Systems), and 2 μM forskolin (Stemcell Technologies), with medium changes every 2–3 days. After predegeneration, tissue was mechanically dissociated with a No. 10 scalpel, digested with 2.5 mg/mL collagenase type I for 30 minutes at 37°C followed by 0.25% trypsin for 10 minutes, treated with DNase I, triturated with a p1000 tip followed by 18-gauge and 21-gauge hypodermic needles, filtered through a 70-μm cell strainer (Nalgene), and plated on poly-L-lysine (Sigma-Aldrich)–coated dishes in Schwann Cell Medium (ScienCell Research Laboratories). Cells were expanded for 6–7 days and cryopreserved in CryoStor CS10 (Stemcell Technologies).

### Fura Red flow cytometry

Cells were thawed, plated on poly-L-lysine–coated plates, and cultured for 7 days. Cells were detached with Accutase (Gibco), blocked with TruStain FcX (BioLegend), and stained with anti-Thy1.2–Alexa Fluor 700 (BioLegend) and Live/Dead Fixable Scarlet (Invitrogen). Cells were loaded with 1 μM Fura Red AM (Invitrogen) and 0.01% Pluronic F-127 (Invitrogen) in HBSS containing Ca^2+^ and Mg^2+^ (Gibco), 0.1% BSA (Sigma-Aldrich), and 10 mM HEPES (Gibco) for 30 minutes at 37°C. After washing, cells were warmed to 37°C for 5 minutes before acquisition on a Cytek Aurora flow cytometer. For stimulated samples, ATP (Sigma-Aldrich) was added to a final concentration of 100 μM 2 minutes before flow cytometry data acquisition. Data were analyzed in FlowJo 10.10.1 after sequential gating on forward and side scatter, singlets, live cells, and Thy1.2-negative cells. A 405-excited/488-excited Fura Red fluorescence ratio was used as a relative calcium-dependent readout.

### Statistics

Data were collated in Microsoft Excel (version 16.114; Microsoft Corp.) and transferred to GraphPad Prism (version 10.6.1; GraphPad Software) for all statistical analyses. Comparisons between two groups were performed using a two-tailed Welch’s t test, which does not assume equal variances between groups. For experiments in which multiple technical or subsampled measurements were obtained from each biological replicate (e.g., g-ratio and circularity measurements), a 2-tailed nested t test was used to account for the hierarchical structure of the data. The specific test used for each experiment is indicated in the corresponding figure legend. Data are presented as mean ± SD, and n represents the number of animals, nerves, or cell batches as noted in the figure legend. A p value of less than 0.05 was considered statistically significant.

### Study approval

All mouse experiments were approved by the University of Pittsburgh Institutional Animal Care and Use Committee (IACUC) under protocol numbers 22061237 and 25066895. Studies were conducted in accordance with the Guide for the Care and Use of Laboratory Animals (National Research Council). The University of Pittsburgh animal care and use program is fully accredited by AAALAC International.

## Data availability

Raw data from RNA-sequencing experiments will be available in the NCBI’s Gene Expression Omnibus database (GEO) under Accession GSE351764 after the October 29, 2026 release date.

## Manuscript information

### Author contributions

JW and DUK conceived and designed the study. JW, MK, MH, AM, OA, and MAJ performed experiments and acquired data. JW, GV, UBP, and DUK supervised experimental work and provided scientific oversight. JW and DUK wrote the manuscript. All authors reviewed and approved the final version of the manuscript.

### AI disclosures

The authors used Claude Opus 5.5 (Anthropic) and GPT-6.1 (OpenAI) to assist in identifying relevant references and to proofread the manuscript for spelling, grammatical, and typographical errors (October 20-26, 2026). All references suggested by the AI tool were independently verified by the authors for accuracy and relevance. The AI tool was not used to generate scientific content, analyze or interpret data, or create figures. The authors reviewed all AI-assisted edits and take full responsibility for the content of the manuscript.

### Conflict of interest

The authors declare no competing financial interests or other conflicts of interest related to this work. No author has received compensation, equity, or other financial benefit from any entity that could be perceived as influencing the research or its reporting. The views expressed in this article are those of the author(s) and do not necessarily reflect the official policy or position of the U.S. Food and Drug Administration (FDA) or the U.S. Department of Health and Human Services. The mention of commercial products, their sources, or their use in connection with material reported herein is not to be construed as either an actual or implied endorsement of such products by FDA.

### Funding support

This work was supported by a Pediatric Scientist Development Program fellowship (NIH 5K12HD000850 subcontract) and the Shear Scholar Award from the Shear Family Foundation (to J.W.); a Richard King Mellon Foundation Institute for Pediatric Research trainee grant and an AFM-Téléthon postdoctoral fellowship (24997) (to O.A.); development funds from the Department of Pediatrics, University of Pittsburgh School of Medicine, an AFM-Telethon grant (23136), Cure CMD research grant, and the NIH Director’s New Innovator Award (DP2-AR081047) (to D.U.K.); and NIH grant R01-AR078872 (initially awarded to D.U.K. and subsequently transferred to G.V.).

## Acknowledgments

We thank members of the Vockley, Pandey, and Kemaladewi laboratories for critical feedback and technical assistance. We also thank Drs. Donna Stolz and Simon Watkins at the University of Pittsburgh Center for Biological Imaging for assistance with electron microscopy, performed using a Leica Ultracut 7 and a JEOL 1400 Plus transmission electron microscope supported by NIH grants 1S10RR025488-01 and 1S10RR016236-01, respectively. We thank Dr. Krishna Prasadan (Imaging Core, Department of Pediatrics, University of Pittsburgh School of Medicine, RRID:SCR_025217) for assistance with confocal imaging; Dr. Radha Gopal, Joshua Michel, and Shushen Xu (Flow Cytometry Core, Department of Pediatrics, University of Pittsburgh School of Medicine, RRID:SCR_025223) for assistance with flow cytometry; Dr. Sunder Sims-Lucas and Riley Gedrys (Histology Core, Department of Pediatrics, University of Pittsburgh School of Medicine, RRID:SCR_027957) for assistance with nerve sectioning; and Dr. Amanda C. Poholek and Dr. William A. MacDonald (Health Sciences Sequencing Core, University of Pittsburgh School of Medicine, RRID:SCR_023116) for assistance with RNA sequencing. The core services and instruments used in this project were graciously supported, in part, by the University of Pittsburgh Department of Pediatrics. Qiagen CLC Genomics Workbench v24 and IPA were licensed through the Molecular Biology Information Service of the Health Sciences Library System at the University of Pittsburgh.

## Notes

### Competing Interest Statement

The authors have declared no competing interest.

## References

1. Helbling-Leclerc A, Zhang X, Topaloglu H, Cruaud C, Tesson F, Weissenbach J, et al. Mutations in the laminin alpha 2-chain gene (LAMA2) cause merosin-deficient congenital muscular dystrophy. Nat Genet. 1995;11(2):216–8.

2. Lokken N, Born AP, Duno M, and Vissing J. LAMA2-related myopathy: Frequency among congenital and limb-girdle muscular dystrophies. Muscle Nerve. 2015;52(4):547–53.

3. Graziano A, Bianco F, D’Amico A, Moroni I, Messina S, Bruno C, et al. Prevalence of congenital muscular dystrophy in Italy: a population study. Neurology. 2015;84(9):904–11.

4. Norwood FL, Harling C, Chinnery PF, Eagle M, Bushby K, and Straub V. Prevalence of genetic muscle disease in Northern England: in-depth analysis of a muscle clinic population. Brain. 2009;132(Pt 11):3175–86.

5. Lake NJ, Phua J, Liu W, Moors T, Axon S, and Lek M. Estimating the Prevalence of LAMA2 Congenital Muscular Dystrophy using Population Genetic Databases. J Neuromuscul Dis. 2023;10(3):381–7.

6. Sarkozy A, Foley AR, Zambon AA, Bonnemann CG, and Muntoni F. LAMA2-Related Dystrophies: Clinical Phenotypes, Disease Biomarkers, and Clinical Trial Readiness. Front Mol Neurosci. 2020;13:123.

7. Xiong H, Tan D, Wang S, Song S, Yang H, Gao K, et al. Genotype/phenotype analysis in Chinese laminin-alpha2 deficient congenital muscular dystrophy patients. Clin Genet. 2015;87(3):233–43.

8. Naom I, D’Alessandro M, Sewry CA, Philpot J, Manzur AY, Dubowitz V, et al. Laminin alpha 2-chain gene mutations in two siblings presenting with limb-girdle muscular dystrophy. Neuromuscul Disord. 1998;8(7):495–501.

9. Gavassini BF, Carboni N, Nielsen JE, Danielsen ER, Thomsen C, Svenstrup K, et al. Clinical and molecular characterization of limb-girdle muscular dystrophy due to LAMA2 mutations. Muscle Nerve. 2011;44(5):703–9.

10. Aumailley M, Bruckner-Tuderman L, Carter WG, Deutzmann R, Edgar D, Ekblom P, et al. A simplified laminin nomenclature. Matrix Biol. 2005;24(5):326–32.

11. Durbeej M. Laminins. Cell Tissue Res. 2010;339(1):259–68.

12. Yurchenco PD, McKee KK, Reinhard JR, and Ruegg MA. Laminin-deficient muscular dystrophy: Molecular pathogenesis and structural repair strategies. Matrix Biol. 2018;71–72:174-87.

13. Timpl R, Tisi D, Talts JF, Andac Z, Sasaki T, and Hohenester E. Structure and function of laminin LG modules. Matrix Biol. 2000;19(4):309–17.

14. Talts JF, Andac Z, Gohring W, Brancaccio A, and Timpl R. Binding of the G domains of laminin alpha1 and alpha2 chains and perlecan to heparin, sulfatides, alpha-dystroglycan and several extracellular matrix proteins. EMBO J. 1999;18(4):863–70.

15. Masaki T, Matsumura K, Saito F, Sunada Y, Shimizu T, Yorifuji H, et al. Expression of dystroglycan and laminin-2 in peripheral nerve under axonal degeneration and regeneration. Acta Neuropathol. 2000;99(3):289–95.

16. Leivo I, and Engvall E. Merosin, a protein specific for basement membranes of Schwann cells, striated muscle, and trophoblast, is expressed late in nerve and muscle development. Proc Natl Acad Sci U S A. 1988;85(5):1544–8.

17. Saito Y, Ishiyama A, Saito Y, Komaki H, and Sasaki M. Myelin abnormalities in merosin-deficient congenital muscular dystrophy. Muscle Nerve. 2024;69(1):55–63.

18. Previtali SC, and Zambon AA. LAMA2 Neuropathies: Human Findings and Pathomechanisms From Mouse Models. Front Mol Neurosci. 2020;13:60.

19. Chan SH, Foley AR, Phadke R, Mathew AA, Pitt M, Sewry C, et al. Limb girdle muscular dystrophy due to LAMA2 mutations: diagnostic difficulties due to associated peripheral neuropathy. Neuromuscul Disord. 2014;24(8):677–83.

20. Shorer Z, Philpot J, Muntoni F, Sewry C, and Dubowitz V. Demyelinating peripheral neuropathy in merosin-deficient congenital muscular dystrophy. J Child Neurol. 1995;10(6):472–5.

21. Quijano-Roy S, Renault F, Romero N, Guicheney P, Fardeau M, and Estournet B. EMG and nerve conduction studies in children with congenital muscular dystrophy. Muscle Nerve. 2004;29(2):292–9.

22. Mercuri E, Pennock J, Goodwin F, Sewry C, Cowan F, Dubowitz L, et al. Sequential study of central and peripheral nervous system involvement in an infant with merosin-deficient congenital muscular dystrophy. Neuromuscul Disord. 1996;6(6):425–9.

23. Deodato F, Sabatelli M, Ricci E, Mercuri E, Muntoni F, Sewry C, et al. Hypermyelinating neuropathy, mental retardation and epilepsy in a case of merosin deficiency. Neuromuscul Disord. 2002;12(4):392–8.

24. Di Muzio A, De Angelis MV, Di Fulvio P, Ratti A, Pizzuti A, Stuppia L, et al. Dysmyelinating sensory-motor neuropathy in merosin-deficient congenital muscular dystrophy. Muscle Nerve. 2003;27(4):500–6.

25. Ghidinelli M, Poitelon Y, Shin YK, Ameroso D, Williamson C, Ferri C, et al. Laminin 211 inhibits protein kinase A in Schwann cells to modulate neuregulin 1 type III-driven myelination. PLoS Biol. 2017;15(6):e2001408.

26. Reinhard JR, Porrello E, Lin S, Pelczar P, Previtali SC, and Ruegg MA. Nerve pathology is prevented by linker proteins in mouse models for LAMA2-related muscular dystrophy. PNAS Nexus. 2023;2(4):pgad083.

27. Qin Y, Akbulut T, Mandraju R, Connolly K, Bechill J, Assadian F, et al. Efficient LAMA1 Gene Activation by Epigenome Editing as a Therapeutic Approach for LAMA2-CMD. Hum Gene Ther. 2026;37(5-6):210–22.

28. Reinhard JR, Lin S, McKee KK, Meinen S, Crosson SC, Sury M, et al. Linker proteins restore basement membrane and correct LAMA2-related muscular dystrophy in mice. Sci Transl Med. 2017;9(396).

29. Tan D, Liu Y, Luo H, Shen Q, Long X, Xu L, et al. A novel mouse model for LAMA2-related muscular dystrophy with analysis of molecular pathogenesis and clinical phenotype. Elife. 2025;13.

30. Kuang W, Xu H, Vachon PH, Liu L, Loechel F, Wewer UM, et al. Merosin-deficient congenital muscular dystrophy. Partial genetic correction in two mouse models. J Clin Invest. 1998;102(4):844–52.

31. Gawlik KI, and Durbeej M. A Family of Laminin alpha2 Chain-Deficient Mouse Mutants: Advancing the Research on LAMA2-CMD. Front Mol Neurosci. 2020;13:59.

32. Miyagoe Y, Hanaoka K, Nonaka I, Hayasaka M, Nabeshima Y, Arahata K, et al. Laminin alpha2 chain-null mutant mice by targeted disruption of the Lama2 gene: a new model of merosin (laminin 2)-deficient congenital muscular dystrophy. FEBS Lett. 1997;415(1):33–9.

33. Sunada Y, Bernier SM, Utani A, Yamada Y, and Campbell KP. Identification of a novel mutant transcript of laminin alpha 2 chain gene responsible for muscular dystrophy and dysmyelination in dy2J mice. Hum Mol Genet. 1995;4(6):1055–61.

34. Pasteuning-Vuhman S, Putker K, Tanganyika-de Winter CL, Boertje-van der Meulen JW, van Vliet L, Overzier M, et al. Natural disease history of the dy2J mouse model of laminin alpha2 (merosin)-deficient congenital muscular dystrophy. PLoS One. 2018;13(5):e0197388.

35. Pellegatta M, De Arcangelis A, D’Urso A, Nodari A, Zambroni D, Ghidinelli M, et al. alpha6beta1 and alpha7beta1 integrins are required in Schwann cells to sort axons. J Neurosci. 2013;33(46):17995–8007.

36. Nodari A, Previtali SC, Dati G, Occhi S, Court FA, Colombelli C, et al. Alpha6beta4 integrin and dystroglycan cooperate to stabilize the myelin sheath. J Neurosci. 2008;28(26):6714–9.

37. Yamada H, Shimizu T, Tanaka T, Campbell KP, and Matsumura K. Dystroglycan is a binding protein of laminin and merosin in peripheral nerve. FEBS Lett. 1994;352(1):49–53.

38. Masaki T, Matsumura K, Hirata A, Yamada H, Hase A, Arai K, et al. Expression of dystroglycan and the laminin-alpha 2 chain in the rat peripheral nerve during development. Exp Neurol. 2002;174(1):109–17.

39. Sagane K, Hayakawa K, Kai J, Hirohashi T, Takahashi E, Miyamoto N, et al. Ataxia and peripheral nerve hypomyelination in ADAM22-deficient mice. BMC Neurosci. 2005;6:33.

40. Lee MK, Tuttle JB, Rebhun LI, Cleveland DW, and Frankfurter A. The expression and posttranslational modification of a neuron-specific beta-tubulin isotype during chick embryogenesis. Cell Motil Cytoskeleton. 1990;17(2):118–32.

41. Harty BL, and Monk KR. Unwrapping the unappreciated: recent progress in Remak Schwann cell biology. Curr Opin Neurobiol. 2017;47:131–7.

42. Griffin JW, and Thompson WJ. Biology and pathology of nonmyelinating Schwann cells. Glia. 2008;56(14):1518–31.

43. Meissner G. The structural basis of ryanodine receptor ion channel function. J Gen Physiol. 2017;149(12):1065–89.

44. Catterall WA. Voltage-gated calcium channels. Cold Spring Harb Perspect Biol. 2011;3(8):a003947.

45. Perez CF. On the footsteps of Triadin and its role in skeletal muscle. World J Biol Chem. 2011;2(8):177–83.

46. Woo JS, Jeong SY, Park JH, Choi JH, and Lee EH. Calsequestrin: a well-known but curious protein in skeletal muscle. Exp Mol Med. 2020;52(12):1908–25.

47. Aguayo-Ortiz R, and Espinoza-Fonseca LM. Linking Biochemical and Structural States of SERCA: Achievements, Challenges, and New Opportunities. Int J Mol Sci. 2020;21(11).

48. Ley K, Laudanna C, Cybulsky MI, and Nourshargh S. Getting to the site of inflammation: the leukocyte adhesion cascade updated. Nat Rev Immunol. 2007;7(9):678–89.

49. Sandor N, Lukacsi S, Ungai-Salanki R, Orgovan N, Szabo B, Horvath R, et al. CD11c/CD18 Dominates Adhesion of Human Monocytes, Macrophages and Dendritic Cells over CD11b/CD18. PLoS One. 2016;11(9):e0163120.

50. Ricklin D, Reis ES, Mastellos DC, Gros P, and Lambris JD. Complement component C3 - The “Swiss Army Knife” of innate immunity and host defense. Immunol Rev. 2016;274(1):33–58.

51. Barratt J, and Weitz I. Complement Factor D as a Strategic Target for Regulating the Alternative Complement Pathway. Front Immunol. 2021;12:712572.

52. Heredia DJ, De Angeli C, Fedi C, and Gould TW. Calcium Signaling in Schwann cells. Neurosci Lett. 2020;729:134959.

53. Ino D, Sagara H, Suzuki J, Kanemaru K, Okubo Y, and Iino M. Neuronal Regulation of Schwann Cell Mitochondrial Ca(2+) Signaling during Myelination. Cell Rep. 2015;12(12):1951–9.

54. Mayer C, Quasthoff S, and Grafe P. Differences in the sensitivity to purinergic stimulation of myelinating and non-myelinating Schwann cells in peripheral human and rat nerve. Glia. 1998;23(4):374–82.

55. Lyons SA, Morell P, and McCarthy KD. Schwann cell ATP-mediated calcium increases in vitro and in situ are dependent on contact with neurons. Glia. 1995;13(1):27–38.

56. Wendt ER, Ferry H, Greaves DR, and Keshav S. Ratiometric analysis of fura red by flow cytometry: a technique for monitoring intracellular calcium flux in primary cell subsets. PLoS One. 2015;10(3):e0119532.

57. Takahashi A, Camacho P, Lechleiter JD, and Herman B. Measurement of intracellular calcium. Physiol Rev. 1999;79(4):1089–125.

58. Kuhlbrodt K, Herbarth B, Sock E, Hermans-Borgmeyer I, and Wegner M. Sox10, a novel transcriptional modulator in glial cells. J Neurosci. 1998;18(1):237–50.

59. Stefansson K, Wollmann RL, and Moore BW. Distribution of S-100 protein outside the central nervous system. Brain Res. 1982;234(2):309–17.

60. Liu Z, Jin YQ, Chen L, Wang Y, Yang X, Cheng J, et al. Specific marker expression and cell state of Schwann cells during culture in vitro. PLoS One. 2015;10(4):e0123278.

61. Brockes JP, Fields KL, and Raff MC. A surface antigenic marker for rat Schwann cells. Nature. 1977;266(5600):364–6.

62. Chang HC, Seki T, Moriuchi T, and Silver J. Isolation and characterization of mouse Thy-1 genomic clones. Proc Natl Acad Sci U S A. 1985;82(11):3819–23.

63. Carenini S, Maurer M, Werner A, Blazyca H, Toyka KV, Schmid CD, et al. The role of macrophages in demyelinating peripheral nervous system of mice heterozygously deficient in p0. J Cell Biol. 2001;152(2):301–8.

64. Kobsar I, Maurer M, Ott T, and Martini R. Macrophage-related demyelination in peripheral nerves of mice deficient in the gap junction protein connexin 32. Neurosci Lett. 2002;320(1-2):17–20.

65. Kaminska J, and Kochanski A. A Role of Inflammation in Charcot-Marie-Tooth Disorders-In a Perspective of Treatment? Int J Mol Sci. 2024;26(1).

66. Osonoi S, Mizukami H, Takeuchi Y, Sugawa H, Ogasawara S, Takaku S, et al. RAGE activation in macrophages and development of experimental diabetic polyneuropathy. JCI Insight. 2022;7(23).

67. Sun W, Yang K, and Song Y. Diabetic neuropathy’s immune-metabolic network: mechanistic complexity, therapeutic challenges, and the path forward. Front Immunol. 2026;17:1804076.

68. Yu YR, O’Koren EG, Hotten DF, Kan MJ, Kopin D, Nelson ER, et al. A Protocol for the Comprehensive Flow Cytometric Analysis of Immune Cells in Normal and Inflamed Murine Non-Lymphoid Tissues. PLoS One. 2016;11(3):e0150606.

69. Newton HS, and Dobrovolskaia MA. Immunophenotyping: Analytical approaches and role in preclinical development of nanomedicines. Adv Drug Deliv Rev. 2022;185:114281.

70. Cossarizza A, Chang HD, Radbruch A, Abrignani S, Addo R, Akdis M, et al. Guidelines for the use of flow cytometry and cell sorting in immunological studies (third edition). Eur J Immunol. 2021;51(12):2708–3145.

71. Saito F, Moore SA, Barresi R, Henry MD, Messing A, Ross-Barta SE, et al. Unique role of dystroglycan in peripheral nerve myelination, nodal structure, and sodium channel stabilization. Neuron. 2003;38(5):747–58.

72. Wright TW, Glowczewskie F, Wheeler D, Miller G, and Cowin D. Excursion and strain of the median nerve. J Bone Joint Surg Am. 1996;78(12):1897–903.

73. Phillips JB, Smit X, De Zoysa N, Afoke A, and Brown RA. Peripheral nerves in the rat exhibit localized heterogeneity of tensile properties during limb movement. J Physiol. 2004;557(Pt 3):879–87.

74. Rosso G, Liashkovich I, Gess B, Young P, Kun A, and Shahin V. Unravelling crucial biomechanical resilience of myelinated peripheral nerve fibres provided by the Schwann cell basal lamina and PMP22. Sci Rep. 2014;4: 7286.

75. Rosso G, Young P, and Shahin V. Implications of Schwann Cells Biomechanics and Mechanosensitivity for Peripheral Nervous System Physiology and Pathophysiology. Front Mol Neurosci. 2017;10:345.

76. Feltri ML, Poitelon Y, and Previtali SC. How Schwann Cells Sort Axons: New Concepts. Neuroscientist. 2016;22(3):252–65.

77. Feltri ML, Graus Porta D, Previtali SC, Nodari A, Migliavacca B, Cassetti A, et al. Conditional disruption of beta 1 integrin in Schwann cells impedes interactions with axons. J Cell Biol. 2002;156(1):199–209.

78. Lee SM, Sha D, Mohammed AA, Asress S, Glass JD, Chin LS, et al. Motor and sensory neuropathy due to myelin infolding and paranodal damage in a transgenic mouse model of Charcot-Marie-Tooth disease type 1C. Hum Mol Genet. 2013;22(9):1755–70.

79. Sadler KE, Lewis TR, Waltz TB, Besharse JC, and Stucky CL. Peripheral nerve pathology in sickle cell disease mice. Pain Rep. 2019;4(4):e765.

80. Guo L, Moon C, Zheng Y, and Ratner N. Cdc42 regulates Schwann cell radial sorting and myelin sheath folding through NF2/merlin-dependent and independent signaling. Glia. 2013;61(11):1906–21.

81. McGowan TJ, Reinhard JR, Lewerenz N, Bialobrzeska M, Lin S, Stepniewski J, et al. Loss of cell-autonomously secreted laminin-alpha2 drives muscle stem cell dysfunction in LAMA2-related muscular dystrophy. Nat Commun. 2025;16(1):10674.

82. Xuan W, and Yao Y. Laminin Receptors in Peripheral Tissues: Functions Revealed by Analysis of Knockout Mice. Am J Pathol. 2025;195(12):2303–19.

83. Sjaastad MD, and Nelson WJ. Integrin-mediated calcium signaling and regulation of cell adhesion by intracellular calcium. Bioessays. 1997;19(1):47–55.

84. Guan JL. Focal adhesion kinase in integrin signaling. Matrix Biol. 1997;16(4):195–200.

85. Yang KJ, Yan Y, Zhang LL, Agresti MA, Matloub HS, LoGiudice JA, et al. Increasing Calcium Level Limits Schwann Cell Numbers In Vitro following Peripheral Nerve Injury. J Reconstr Microsurg. 2017;33(6):435–40.

86. Yi B, Zhang K, Chen X, Xu R, Zhang X, Zhang C, et al. Bioinspired aligned electroconductive hydrogel nanofiber patch enhances peripheral nerve repair through mechanosensitive calcium influx and focal adhesion kinase/protein kinase B pathway activation. Acta Biomater. 2026;217:243–57.

87. Corral JG, Cheli VT, Santiago-Gonzalez DA, Kedari KG, and Paez PM. Chemogenetic Schwann cell activation impairs early myelination and triggers adult demyelination in the peripheral nervous system. Front Cell Neurosci. 2026;20:1771951.

88. Yuan Y, Wang Y, Wu S, and Zhao MY. Review: Myelin clearance is critical for regeneration after peripheral nerve injury. Front Neurol. 2022;13:908148.

89. Lindborg JA, Mack M, and Zigmond RE. Neutrophils Are Critical for Myelin Removal in a Peripheral Nerve Injury Model of Wallerian Degeneration. J Neurosci. 2017;37(43):10258–77.

90. Liu P, Peng J, Han GH, Ding X, Wei S, Gao G, et al. Role of macrophages in peripheral nerve injury and repair. Neural Regen Res. 2019;14(8):1335–42.

91. de Laat ECM, Groothuis JT, Bouman K, Houwen-van Opstal SLS, Erasmus CE, and Voermans NC. Clinical and functional outcome measures in LAMA2-related muscular dystrophy and SELENON-related myopathy; a 1.5-year natural history study. J Neuromuscul Dis. 2026:22143602261465467.

92. Hinkley L, Orbach R, Park J, Alvarez R, Dziewczapolski G, Bonnemann CG, et al. An international retrospective early natural history study of LAMA2-related dystrophies. J Neuromuscul Dis. 2024;11(6):1238–46.

93. de Laat ECM, Houwen-van Opstal SLS, Bouman K, van Doorn JLM, Cameron D, van Alfen N, et al. A 5-year natural history study in LAMA2-related muscular dystrophy and SELENON-related myopathy: the Extended LAST STRONG study. BMC Neurol. 2024;24(1):409.

94. Camelo CG, Artilheiro MC, Fernandes TR, Moreno CAM, Fonseca A, Reed UC, et al. Cross-sectional survey study of the natural history of LAMA2-related dystrophy. Clin Neurol Neurosurg. 2024;245:108467.

95. Enzmann C, Steiner L, Pospieszny K, Zweier C, Plattner K, Baumann D, et al. A Multicenter Cross-Sectional Study of the Swiss Cohort of LAMA2-Related Muscular Dystrophy. J Neuromuscul Dis. 2024;11(5):1021–33.

96. Kemaladewi DU, Bassi PS, Erwood S, Al-Basha D, Gawlik KI, Lindsay K, et al. A mutation-independent approach for muscular dystrophy via upregulation of a modifier gene. Nature. 2019;572(7767):125–30.

97. Rafel-Millan N, Bayod-Giron C, Pallares-Masmitja M, Fernandez-Aviles C, Figuereo-Moya K, Edo A, et al. Triple-AAV intein-mediated gene therapy ameliorates dystrophic phenotype in MDC1A mice. Mol Ther. 2026;34(10):5697–709.

98. Kaiser T, Allen HM, Kwon O, Barak B, Wang J, He Z, et al. MyelTracer: A Semi-Automated Software for Myelin g-Ratio Quantification. eNeuro. 2021;8(4).

