## Supplemental Figures for "LAMA2-related muscular dystrophy alters Schwann cell calcium homeostasis and increases peripheral nerve inflammation"

**Supplemental figures and legends**

Supplemental Figure 1: LAMA2 protein localizes to the periphery of Remak bundles. Representative image of sciatic nerves from 25-week-old wild-type C57BL/6J mice immunostained against the indicated antigens. Arrows indicate Remak bundles (bundles of axons surrounded by the cytoplasm of a non-myelinating Remak Schwann cell). LAMA2 signal is present around the outer edge of each Remak bundle and is not detectable at the interface between axon and Remak Schwann cell. Sciatic nerves from 3 mice were assessed. Scale bar = 5 µm.

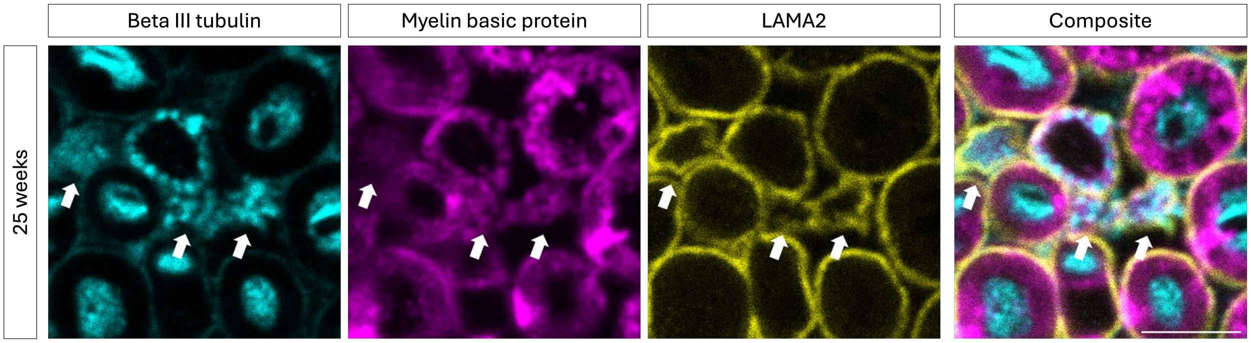

Supplemental Figure 2: Secondary antibody only controls for immunostaining.

A and B) Sciatic nerve immunostaining with secondary antibody only from animals of the indicated ages. Scale bar = 10 μm. Images are scaled identically to corresponding immunostaining in Figure 1, B. C and D) Sciatic nerve immunostaining with secondary antibody only from animals of the indicated ages. Scale bar = 100 μm. Images are scaled identically to corresponding immunostaining in Figure 5, A and C. Nerves from 3-5 mice were assessed for each condition.

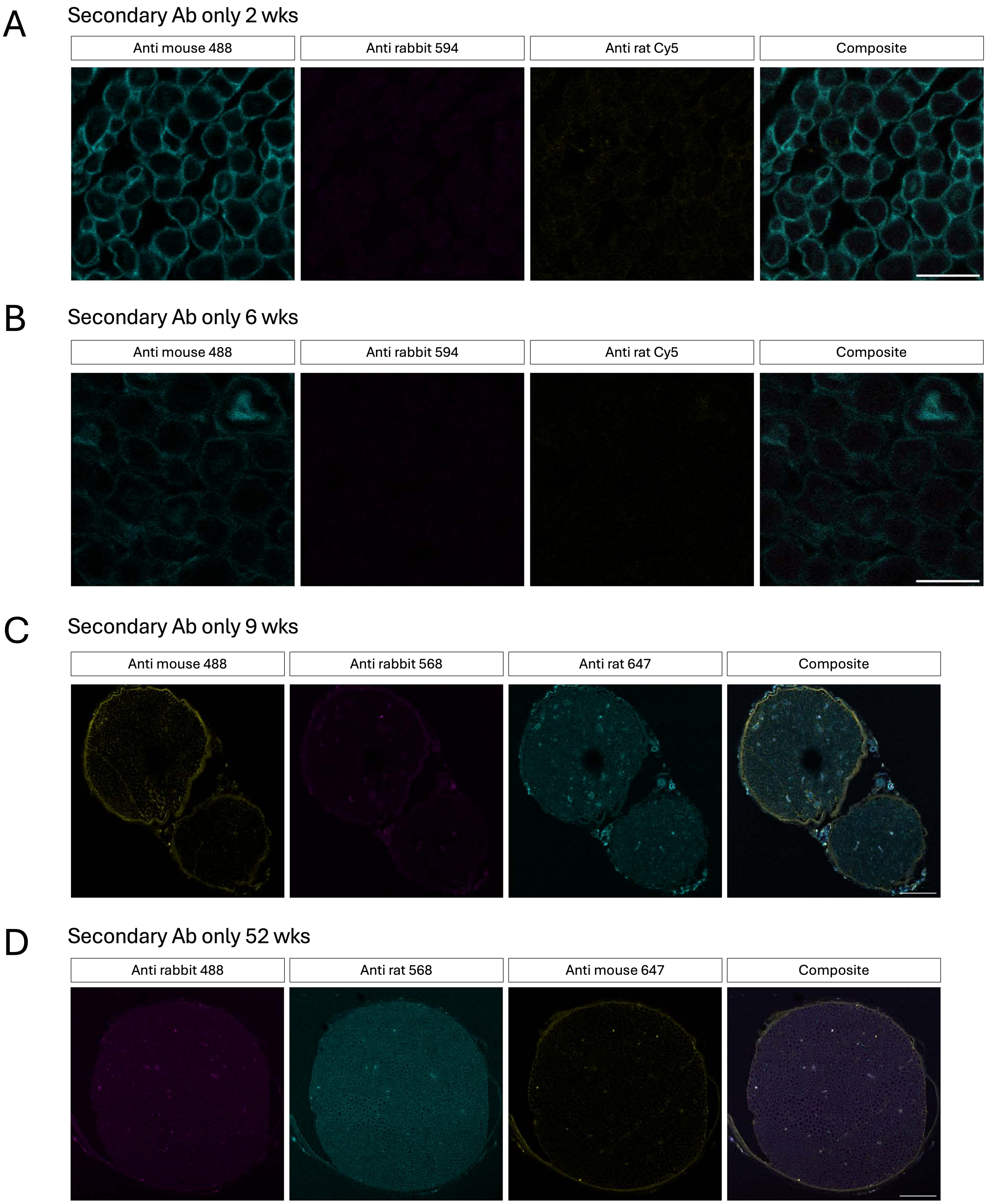
A
Secondary Ab only 2 wks
Anti mouse 488
Anti rabbit 594
Anti rat Cy5
Composite
B
Secondary Ab only 6 wks
Anti mouse 488
Anti rabbit 594
Anti rat Cy5
Composite
C
Secondary Ab only 9 wks
Anti mouse 488
Anti rabbit 568
Anti rat 647
Composite
D
Secondary Ab only 52 wks
Anti rabbit 488
Anti rat 568
Anti mouse 647
Composite

Supplemental Figure 3: Transmission electron microscopy of unmyelinated regions in wild-type and LAMA2-deficient sciatic nerves.

Transmission electron micrograph of A) Remak bundles in wild-type sciatic nerves, B) Remak bundles in LAMA2-deficient sciatic nerves, and C) unmyelinated naked axons in LAMA2-deficient sciatic nerves taken at 25,000× magnification. Arrowheads indicate a representative pair of double membranes of adjacent axons in Remak bundles. Arrows indicate a representative pair of single membranes in adjacent naked unmyelinated axons. Nerves from 4-5 mice were assessed for each condition. Scale bar = 1,000 nm.

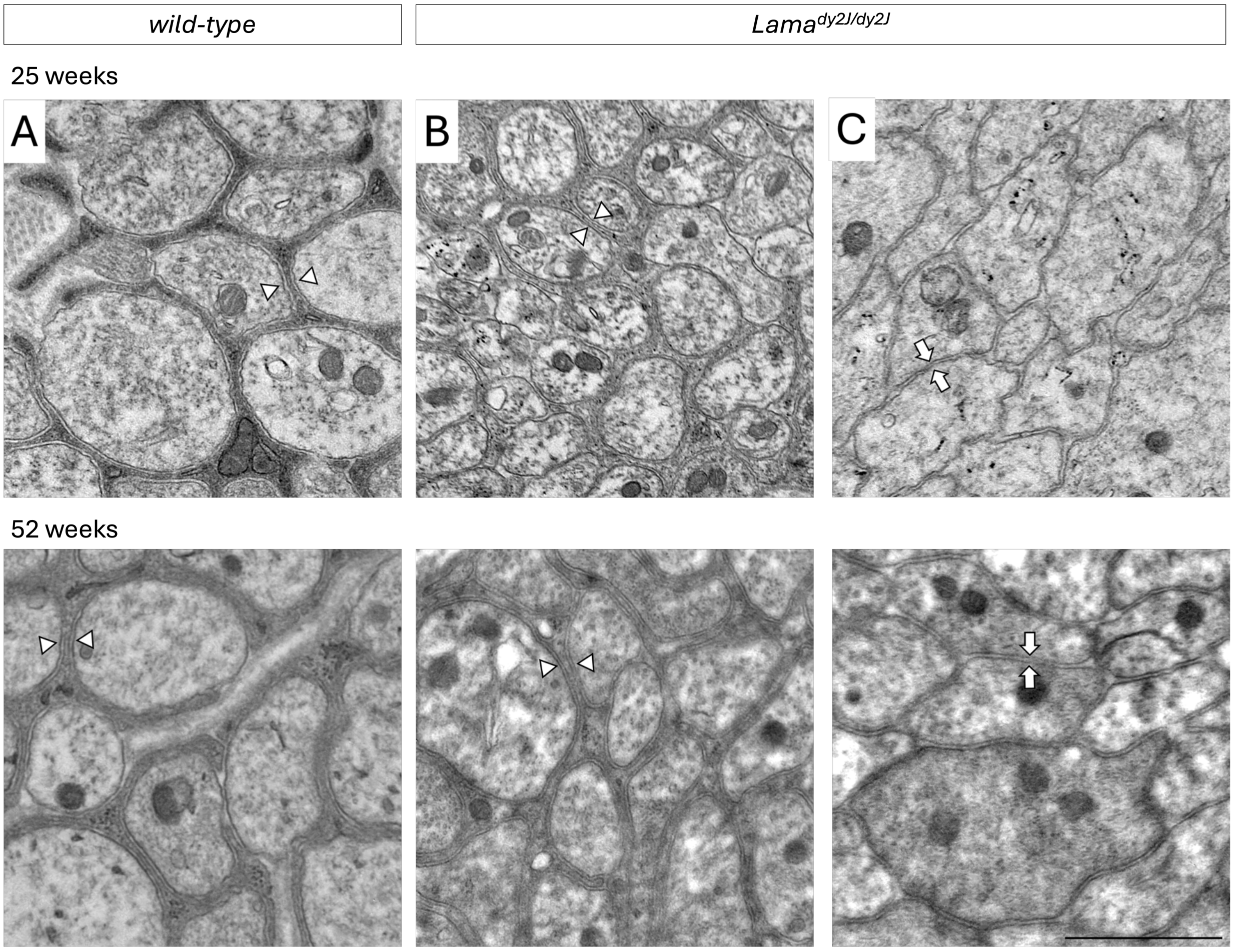

Supplemental Figure 4: Other immune-related pathways are mostly upregulated in LAMA2-deficient nerves.

Predicted activation z-score of the A) pathogen-induced cytokine storm signaling pathway and B) the phagosome formation pathway at the indicated timepoints. Sequencing data from 4-6 mice of each genotype and age were analyzed.

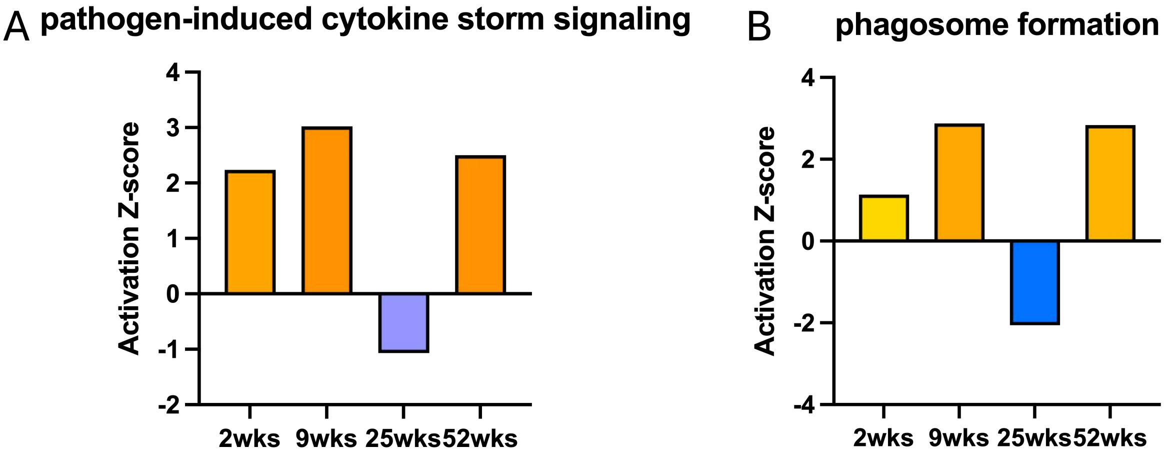

Supplemental Figure 5: Calcium signaling is downregulated and immune-related pathways are upregulated in LAMA2-null nerves.

Volcano plot of signaling pathways predicted to be altered based on RNA sequencing data at 2 weeks in LAMA2-dyW nerves compared to wild-type controls. Two wild-type and three LAMA2-dyW nerves were analyzed. X axis represents activation z-score, with negative values indicated by blue bubbles and positive values indicated by orange bubbles. Y axis represents statistical significance, with high values representing increased statistical significance. The size of each bubble corresponds to the number of differentially expressed genes mapped to that pathway.

Alteration of signaling pathways in *Lama2*$^{dyW/dyW}$ sciatic nerves at 2 weeks

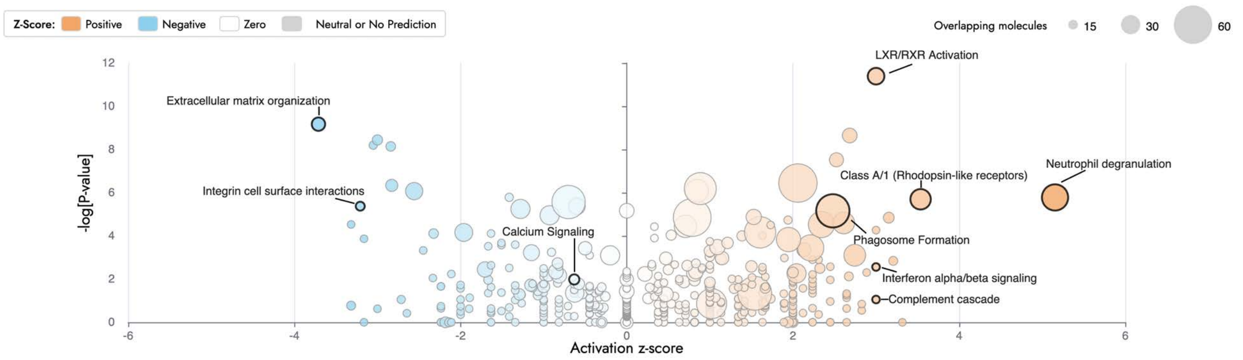

Supplemental Figure 6: Simplified schematic of calcium homeostasis and calcium-dependent processes.

Simplified schematic of components of calcium homeostasis and calcium-dependent processes. The top differentially expressed genes and processes at 25 weeks identified by RNA sequencing in Figure 3, D are indicated in red.

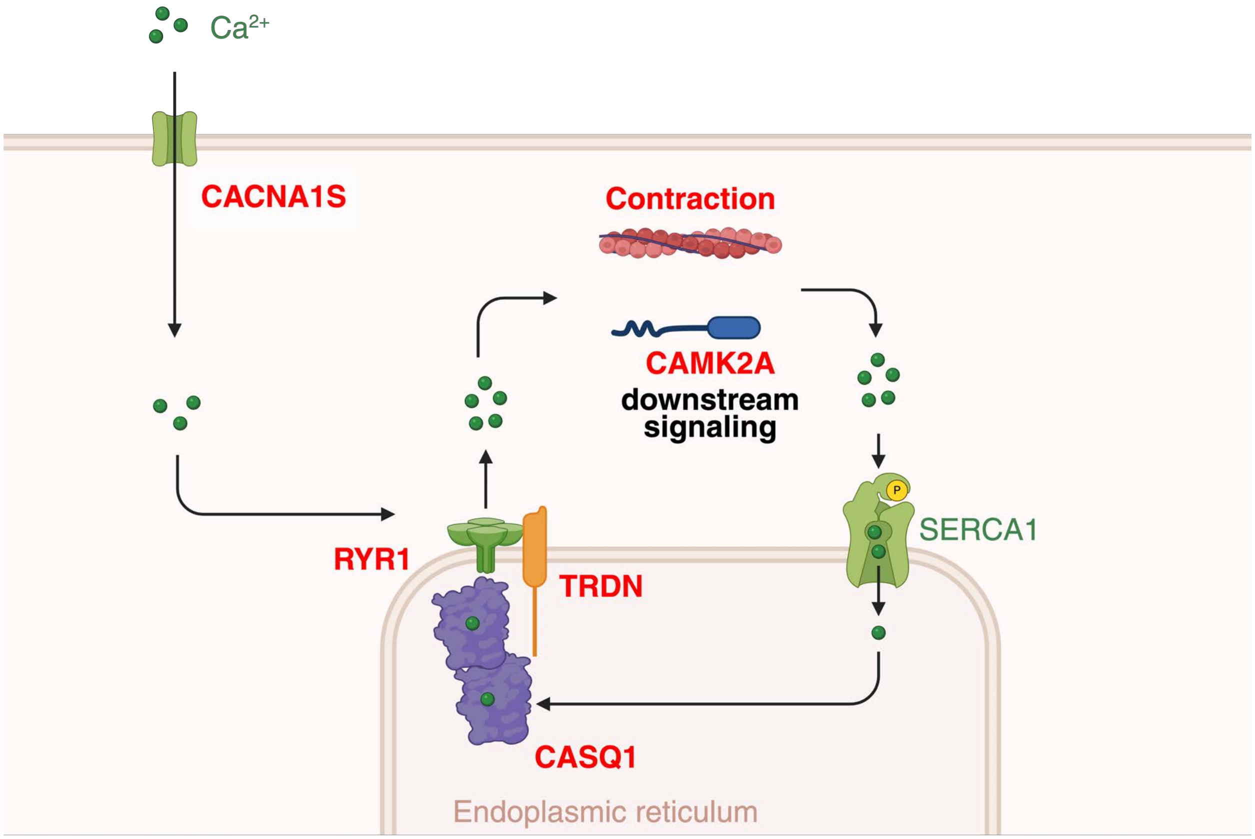

Supplemental Figure 7: Marker immunostaining of primary Schwann cells.

A) Schematic of primary Schwann cell culture derivation. B) Immunostaining of primary Schwann cell cultures against Schwann cell markers S100B (yellow) and Sox10 (magenta). Cultures are a mixture of Schwann cells (S100B$^{+}$ and Sox10$^{+}$) and other cells. Scale bar = 100 μm.

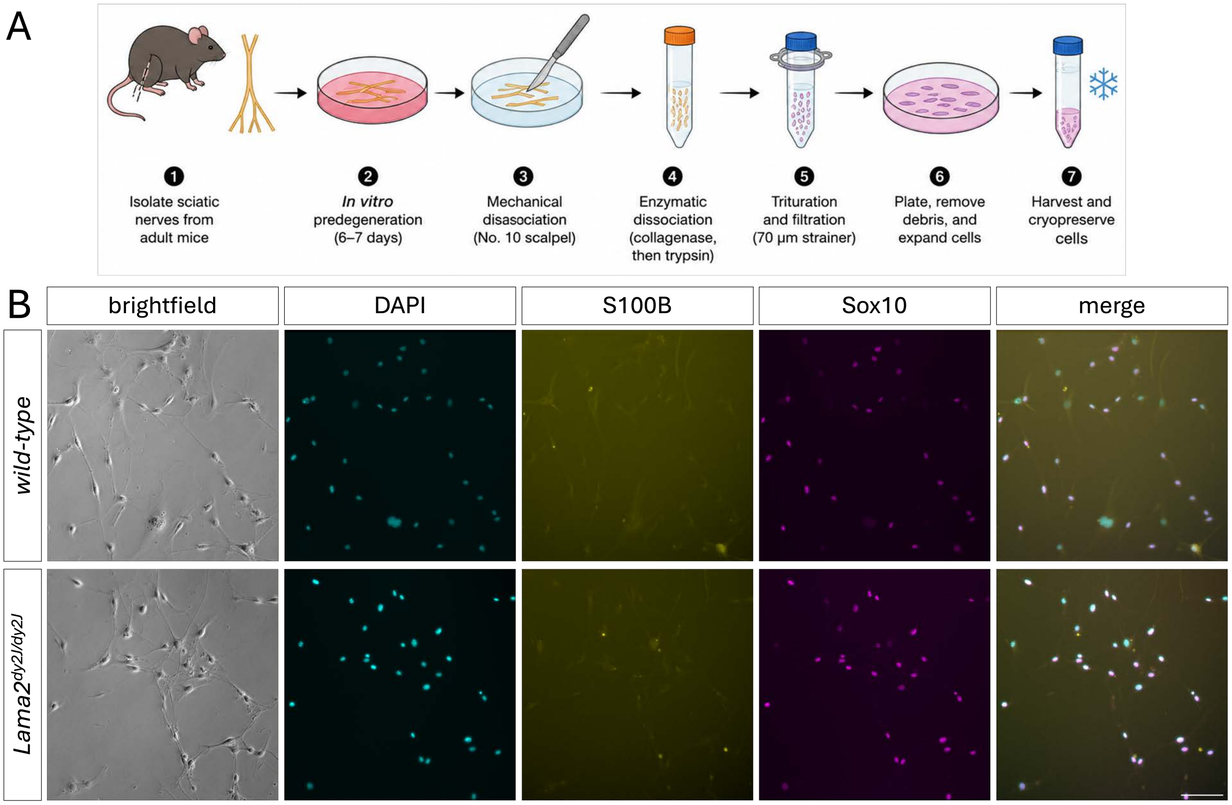

Supplemental Figure 8: Gating strategy for flow cytometry.

Cells were gated on forward and side scatter. Singlets were selected. Thy1.2-A700 negative and Live/Dead Scarlet negative cells were selected for 405:488 fluorescence ratio measurements, with gating based on unstained and single-positive cell plots.

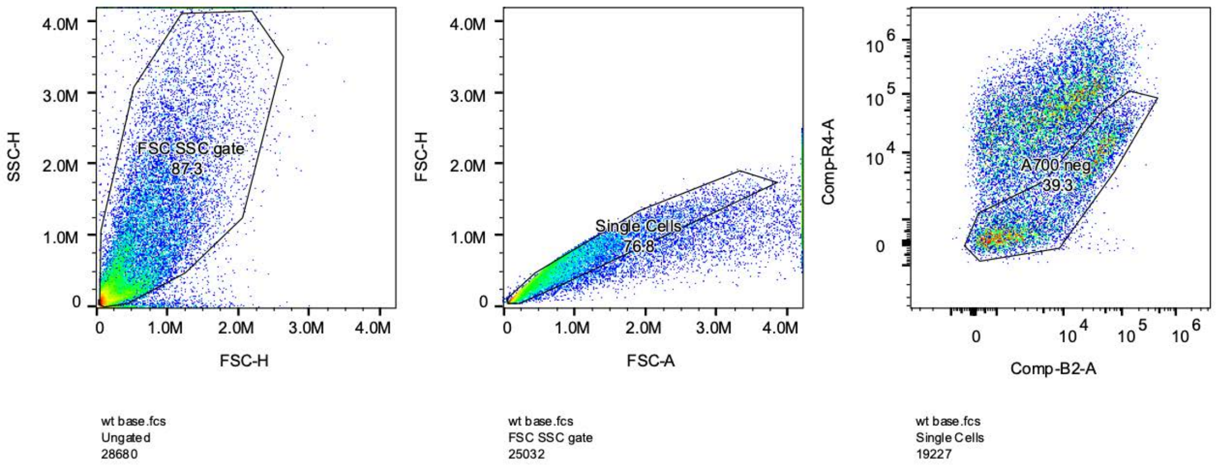
